# Cyanopeptides Restriction and Degradation Co-mediate Microbiota Assembly During a Freshwater Cyanobacterial Harmful Algal Bloom (CyanoHAB)

**DOI:** 10.1101/2021.12.31.474673

**Authors:** Han Gao, Ze Zhao, Lu Zhang, Feng Ju

## Abstract

Cyanobacterial harmful algal blooms (CyanoHABs) are globally intensifying and exacerbated by climate change and eutrophication. However, microbiota assembly mechanisms underlying CyanoHABs remain scenario specific and elusive. Especially, cyanopeptides, as a group of bioactive secondary metabolites of cyanobacteria, could affect microbiota assembly and ecosystem function. Here, the trajectory of cyanopeptides were followed and linked to microbiota during *Microcystis*-dominated CyanoHABs in lake Taihu, China. The most abundant cyanopeptide classes detected included microginin, spumigin, microcystin, nodularin and cyanopeptolin with total MC-LR-equivalent concentrations between 0.23 and 2051.54 ppb, of which cyanotoxins beyond microcystins (e.g., cyanostatin B and nodularin_R etc.) far exceeded reported organismal IC_50_ and negatively correlated with microbiota diversity, exerting potential collective eco-toxicities stronger than microcystins alone. The microbial communities were differentiated by size fraction and sampling date throughout CyanoHABs, and surprisingly, their variances were better explained by cyanopeptides (19-38%) than nutrients (0-16%). Cyanopeptides restriction (e.g., inhibition) and degradation are first quantitatively verified as the deterministic drivers governing community assembly, with stochastic processes being mediated by interplay between cyanopeptide dynamics and lake microbiota. This study presents an emerging paradigm in which cyanopeptides restriction and degradation co-mediate lake water microbiota assembly, unveiling new insights about the ecotoxicological significance of CyanoHABs to freshwater ecosystems.

## 1. Introduction

Cyanobacterial harmful algal blooms (CyanoHABs) increasingly threaten marine and freshwater ecosystems worldwide (Huisman et al. 2018, Wang et al. 2021), as cyanobacteria produce diverse bioactive non-ribosomal oligopeptide metabolites known as cyanopeptides (e.g., microcystins, nodularins, spumigins, anabaenopeptins, cyanopeptolins, microginins, microviridins and aerucyclamides) (Janssen 2019). A few recent studies have demonstrated the largely neglected ecological impacts (abbreviated as ‘eco-impacts’, e.g., on nutrients and carbon turnover (Egli et al. 2020)) or life toxicities (e.g., for larval fish (Fernandes et al. 2019)) caused by non-microcystin cyanopeptides, such as microginin, cyanopeptolin A and arucyclamide A (Beversdorf et al. 2018, Fernandes et al. 2019, Huang and Zimba 2019). Meanwhile, non-microcystin cyanopeptides have been detected at significant amounts in the surface drinking water sources adversely affected by CyanoHABs, suggesting their ineffective removal by drinking water treatment plants and potential threats to human health (Beversdorf et al. 2018). Still, compared to the microcystins, non-microcystin cyanopeptides, especially for their dynamics in environments and eco-impacts, received much less attention. Exploring cyanopeptides beyond microcystins is critical for elucidating the eco-impacts and health risk posed by CyanoHABs in freshwater ecosystems, particularly for those serving as drinking water reservoirs.

Taihu, a shallow, subtropical lake (an average depth of 1.9 m), is the source of drinking and irrigating water for more than 40 million residents around its watershed (Qin et al. 2019). Extensive urbanization and hydrological engineering (e.g. Project of Water Diversion from the Yangtze River to Lake) over the decades have bred the eutrophic and fluctuating status of its water quality, exacerbating the prevalence of CyanoHABs (Huang et al. 2021, Huang et al. 2019, Qin et al. 2019, Qin et al. 2020), of which the causative taxa include prolific cyanopeptide producers like *Microcystis* and *Dolichospermum* (Ma et al. 2016, Osterholm et al. 2020). Current knowledge on cyanopeptide in Taihu is primarily constricted to microcystins (e.g., MC-LR, MC-RR and MC-YR) in water (Sakai et al. 2013, Su et al. 2018, Tang et al. 2018, Xue et al. 2020), sediment (Xue et al. 2020), and nearshore soil (Chen et al. 2006). In contrast, non-microcystin components and their dynamics have not been characterized in the changing hydrological and ecological conditions. Untangling the spatiotemporal dynamics and eco-impacts of those CyanoHABs byproducts can provide basis for their health risk assessment, and further inform management of water withdraw or water diversion to prevent their detrimental effects.

Microbiota underlying CyanoHABs include cyanopeptide producers (i.e., sources) and degraders (i.e., sinks), and their spatiotemporal assemblages are closely linked with cyanopeptide dynamics. Specifically, taxonomic composition and biogeochemical function of bloom-associated microbiota showed clear succession over the course of toxigenic blooms (Tang et al. 2018, Wan et al. 2019). Size fractions of biomass can form distinct ecological niches within which interactions between cyanobacteria and bloom-associated microbiota are imperative and diverse (Chen et al. 2018, Cook et al. 2020, Tang et al. 2017, Xu et al. 2018, Zhu et al. 2021a). For example, microorganisms colonized on cyanobacterial aggregates (> 100 μm) are functionally linked with their host in a commensal relationship (Cook et al. 2020). Although taxonomic overlap exists between heterotrophic bacteria from particle-associated (> 5 μm) and free-living fractions (2-0.2 μm) (Tang et al. 2017), their community functions and environmental preference are different (Xu et al. 2018). However, the relative contribution of cyanopeptides to the assembly of the niche-differentiated microbiota during CyanoHABs has not been evaluated. Considering the combined effects of their antimicrobial activities (Swain et al. 2017) and nutritional niches during degradation (Li et al. 2017), a much more complicated scenario than previously appreciated may exist regarding niche-based deterministic processes (e.g., differentiation) and their role in mediating assembly processes of bloom water microbiota. A fundamental but unresolved question arises: How could dynamics of these bioactive metabolites, as a selective driver, shape cyanobacterial and non-cyanobacterial communities across size-fractionated bloom water microbiota? Unravelling ecological rules guiding microbiota assembly during bloom events, especially the interactions between cyanopeptides and microbiota should provide a theoretical guidance for microbiome-based control of CyanoHABs byproducts.

Using lake Taihu as a model system, this study aims to i) resolve spatiotemporal dynamics of cyanopeptides and their producers over the course of CyanoHABs; ii) explore how the degradation of cyanopeptides is related to documented degraders or microbial setups; and 3) explore how and to what extent can the cyanopeptides affect microbiota assemblages in different size fractions of bloom-associated community. Keeping those in mind, we carried out a series of field sampling campaigns in three lake regions in northeast Taihu and set up in-lab microbial degradation assays. A list of cyanopeptides was identified through non-targeted screening of cyanobacterial extracts using high resolution mass spectrometer (HR-MS) based on a recently published cyanopeptide database (Jones et al. 2021), and then quantified by tandem mass spectrometer (MS/MS). Moreover, 16S rRNA gene-based absolute abundance quantification enables a quantitative view on the contrasting changes in the biomass and diversity of microbiota, and further linking to cyanopeptide dynamics. Examining the predominant cyanopeptides and associated microbiota would not only advance current understanding of their yet-to-be-recognized eco-impacts on the assembly of lake water microbiota, but also provide guidance for ecotoxicological risk assessment and management policy of CyanoHABs in freshwater ecosystem.

## 2. Methods

### 2.1 Sample collection

Water samples were collected between June 9^th^ and September 20^th^, 2020 on five dates to cover a cyanobacterial bloom process. Eight sites (shown in Fig. 1A) were selected representing three lake regions: the Meiliang Bay (Site 5 and 6), Gonghu Bay (Site 1, 2, and 16) and the open water connecting the two bays (Site 3, 4 and 8). For the surface water (ca. 0-0.4 m of the upper water column), a 48-μm pore size sieve was used to collect cyanobacterial colonies (scum) from 5-liter of surface water by a 2.5-liter organic glass water sampler (JC-800, Juchuang, China). The scum was backwashed into a 50-ml centrifuge tube with a final volume of 40 ml using Milli-Q water. The filtrate was kept in purified water bottles pre-rinsed with the filtrate. For cyanopeptide extraction, both surface and overlaying water (ca. 0-0.4 m above the sediment) samples were preserved in 50-ml centrifuge tubes. Water temperature, pH, conductivity and dissolved oxygen was determined *in situ* by a portable multimeter (HQ40D, HACH, USA). Additionally, 500 ml of lake water was collected using a polycarbonate bottle for analysis of total nitrogen (TN), total phosphorus (TP), dissolved total nitrogen (DTN), dissolved total phosphorus (DTP), phosphate, nitrate and ammonium. And dissolved organic nitrogen (DON) was calculated from difference between TN and inorganic nitrogen (nitrate and ammonium). All the tubes and bottles were stored in a cooler (80 L, CONTOOSE, China) with icebags and delivered back to the lab within 5 hours.

**Figure 1.**
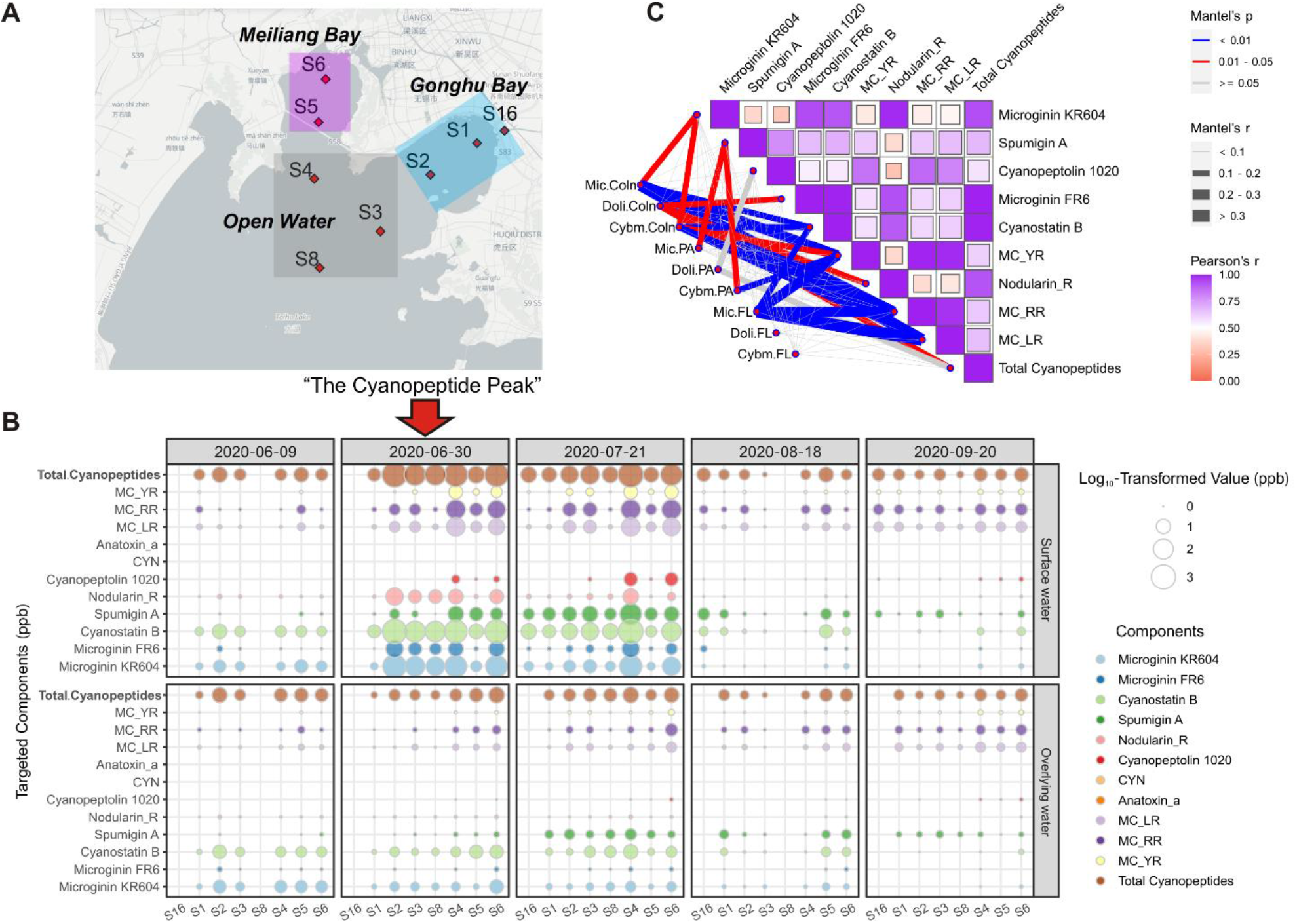
Spatiotemporal heterogeneity of cyanopeptides and its links with cyanobacterial communities. (A) Sampling sites in Taihu. The purple, grey and blue rectangles represent the lake region Meiliang Bay, Open water and Gonghu Bay, respectively. (B) The log_10_-transformed concentration (ppb or µg L^-1^) of cyanopeptides detected in eight sampling sites at five sampling dates & (C) Correlations between dominant cyanobacterial genera and cyanopeptide compounds computed by Mantel test. The widths of lines denoted the value of Mantel’s r. Colors of edges are distinguished based on P values. Pearson correlation coefficients between cyanopeptide components were visualized in the heatmap. The results showed that cyanopeptides in the lake water was dominated by components from the microginins, microcystins and spumigins class, reaching a cyanopeptide peak on June 30. Significant correlations of cyanopeptide components with those dominant cyanobacterial genera were observed, especially within the colony fraction. Abbreviations used in the figures are, Cyano: Cyanobacteria, Coln: colony, PA: particle-associated, FL: free-living, Mic: *Microcystis*, Doli: *Dolichospermum* and Cybm: *Cyanobium*.

### 2.2 DNA extraction and 16S rRNA gene analysis

To obtain size-fractioned microbial DNA, 250 to 300 ml pre-sieved lake water was sequentially vacuum filtered with 2.0 μm and 0.2 μm pore size polycarbonate filters (Millipore, USA). Then, DNA extraction was performed using DNeasy Powersoil Kit (Qiagen, Germany) following the manufacture’s instruction (see Supporting Information S1 for details). To enable absolute quantification of the microbial communities, 16S rRNA gene copy number of each DNA extract was determined by quantitative PCR assay as described in (Ju et al. 2019) and also detailed in Supporting Information S1. The V4-V5 regions of prokaryotic 16S rRNA gene were targeted by the primers F515 (GTGYCAGCMGCCGCGGTAA) and R926 (CCGYCAATTYMTTTRAGTTT). Sequencing libraries were generated by NEBNext ® Ultra™ □ DNA Library Prep Kit for Illumina®. Paired-end sequencing was conducted on an Illumina Nova6000 platform with read length set to 250 base pairs (Guangdong Magigene Biotechnology Co., Ltd. Guangzhou, China).

The raw reads were analyzed using the QIIME2 (Bolyen et al. 2019) followed in-house Python scripts for batch processing (see Supporting Information S2 for details). The data that support the findings of this study have been deposited into CNGB Sequence Archive (CNSA) of China National GeneBank DataBase (CNGBdb: www.cngb.org/cnsa) with accession number CNP0001905 (http://db.cngb.org/cnsa/project/CNP0001905/reviewlink/).

### 2.3 Extraction, identification and quantification of cyanopeptides

Raw water collected from both the surface and bottom (overlaying water) for cyanopeptides quantification were put through two frozen-thaw cycles to release intracellular components. Solid phase extraction (SPE) was used for extraction and purification. A target list of cyanopeptides was first determined by injecting 10 μl concentrated cyanopeptide extracts (collected on July 21) into a high-resolution mass spectrometer (Thermo Orbitrap Fusion Lumos, Thermo, USA). Candidates with at least two daughter ions hit the reference were reserved (Table S1 & Fig. S1), ending up with a list including nine cyanopeptides components. Quantification was then performed on a UPLC-MS/MS system as described in Supporting Information S3.

### 2.4 Microcystin degradation experiments

For two of the sampling campaigns (on June 30 and July 21), in-lab experiments were conducted to assess the capability of lake water in degrading microsystins. Experimental setups were described in Supporting Information S4. To link degradation rate to documented degrading organisms, a list containing bacterial genera reported as organisms capable of degrading microcystin was summarized from published literatures (as summarized in Table S2), which served as a library to retrieve a subset of potential degrading taxa from the sequencing data.

### 2.5 Statistical analyses

For each sample, the total copy numbers of 16S rRNA gene measured by qPCR were multiplied by relative abundances of each ASV. The volume of water filtered or concentrated during sampling was used to compute the absolute abundance of each ASV expressed as ‘copy numbers per liter of lake water (copies L^-1^) and used in downstream statistical analysis (see Supporting Information S5).

To quantify ecological stochasticity of size-fractioned microbial communities, normalized stochasticity ratio (NST) of cyanobacterial or non-cyanobacterial communities in three size fractions were calculated using *NST* package in R. The NST metric is developed as a null-model-based estimation to predict whether community assembly processes are more stochastic or more deterministic (Ning et al. 2019). Fifty percent is considered a boundary where deterministic and stochastic processes are categorized.

## 3. Results

### 3.1 Spatiotemporal distribution of cyanopeptides correlated with cyanobacterial communities

Nine major variants belonging to five cyanopeptide classes including one nodularin, three microginins, one spumigin, and one cyanopeptolin were co-identified (Table S1). The average concentrations of all the nine variants reached a “Cyanopeptide Peak” on June 30 (ca. 597 ppb, n = 7) and declined by 50% on July 21 (ca. 300 ppb, n = 8). The cyanopeptides were primarily accumulated on the surface but can also be detected in the overlying water (Fig. 1B). In surface water, cyanostatin B was detected in water column from below limit of quantification to a highest record of 1323.81 ppb (MC-LR-equivalent concentration) on July 21 at Site 4, two orders of magnitude higher than the synchronously detected MC-LR (52.53 ppb) and MC-RR (91.63 ppb). Besides, the contents of microginin (FR6 and KR604) showed ∼200% higher concentrations than microcystin components (Fig. 1B). Geographically, the highest total concentrations of cyanopeptides were detected in samples from Site 2 (1859.09 ppb) in Gonghu Bay, Site 6 (963.52 ppb) in Meiliang Bay, and Site 4 (2051.54 ppb) in the open water (Fig. 1B).

To profile microbiological producers of major cyanopeptides and their spatiotemporal dynamics, 102 size-fractionated biomass DNA samples were quantitatively analyzed using high-throughput quantitative 16S rRNA gene amplicons sequencing. The results showed a sharp temporal change in both bacterial (Fig. S3) and cyanobacterial (Fig. 2A) biomass over the course of blooming, accompanying a contrasting succession in the cyanobacterial community taxonomic composition (Fig. 2B) and alpha-diversity (Fig. S4A-4C) across the lake regions and size fractions examined. The total cyanobacterial biomass was significantly and continuously elevated until reaching a Biomass Peak on July 21 (Fig. 2A), whereas the alpha diversity metrics (e.g., observed species, Chao 1 richness and Shannon’s H) of cyanobacterial communities significantly decreased (Fig. S4A). This contrasting pattern consisted with the much lower alpha diversity metrics within the colony fractions than PA or FL fractions (*P* < 0.01, Fig. S4C), reflecting the prosperity of bloom-thrived cyanobacterial species at the expense of compressing living conditions of the others. Furthermore, non-metric multidimensional scaling (NMDS) analysis showed that the colony fractions harbored cyanobacterial communities structurally distinct from the FL or PA fractions (Fig. S4). Accordingly, size fraction was more pronounced in explaining community dissimilarities compared to sampling date and lake region (Table 1).

**Figure 2.**
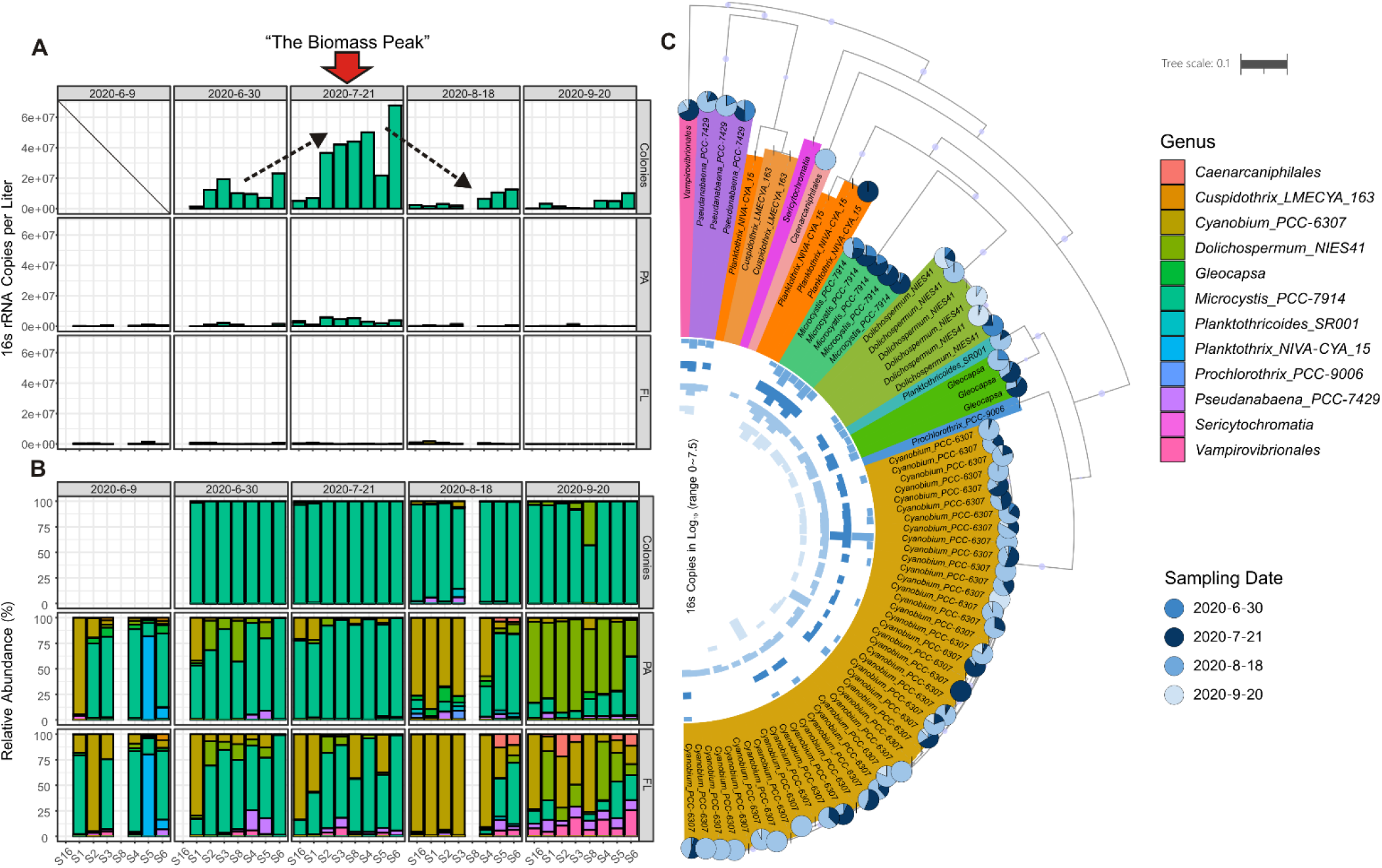
Spatiotemporal dynamics of biomass and composition in three size fractions of the cyanobacterial communities. The analysis was based on 16S rRNA gene sequencing of three fractions of the microbiota defined by water filtration: > 48 μm (colony fraction), 2.0-0.2 μm (particle associated) and < 0.2 μm (free living). (A) 16S-based absolute abundance (copies L^-1^) or biomass estimates of cyanobacterial genera. (B) Relative abundance (%) of cyanobacterial genera. (C) Phylogenetic tree of all cyanobacterial ASVs and abundance distribution analyses of cyanobacterial populations in the colony fraction. Barplots in the inner cycle represent the log-transformed absolute abundance of each ASV on four sampling dates. Pie plots at the tips of each ASV represent it relative proportions on four sampling dates. Results demonstrated a clear biomass peak of cyanobacterial bloom on July 21 and succession of the dominant genera on Aug. 18 and Sept. 20 in PA and FL fractions with high micro-diversity.

Throughout the whole sampling period, *Microcystis* was the most abundant genus (6.9×10^6^ to 6.8×10^7^ copies L^-1^), primarily aggregating as colonies (Fig. 2A). Its species largely died off on the August 18 and September 20 (ca. 83.3% reduction in biomass), accompanying a dramatic increase in the average relative abundance of *Cyanobium* and *Dolichospermum* (from 22.5% to 65.8% in total) in the PA and FL fractions of the biomass (Fig. 2B). To resolve cyanobacterial diversity within the cyanobacterial colonies down to the species or population level, a phylogenetic tree was built using cyanobacterial 16S rRNA gene ASVs (Fig. 2C). We then comparatively examine both the 16S-based absolute abundance (copies L^-1^, bar charts at inner cycle) and relative proportions of each ASV (pies charts) in the colony fraction on four sampling dates (Fig. 2C) and found the co-existence of multiple closely related ASVs within *Microcystis* (5 ASVs) with changing abundances. Similarly, high micro-diversity and population succession dynamics were also observed within genera, e.g., *Cyanobium* spp. (43 ASVs) and *Dolichospermum* spp. (6 ASVs, Fig. 2C).

Mantel tests were performed to explore associations between the 162 ASVs assigned to the major cyanobacterial genera (i.e., *Microcystis, Cyanobium* and *Dolichospermum*, total relative abundance > 90%) and cyanopeptide profiles (Fig. 1C). Generally, microcystins, microginin KR604 and cyanostatin B were correlated significantly (Mantel’s *r* = 0.155 to 0.367, *P* < 0.05) and frequently (n = 10 links) with the abundance matrix of *Microcystis* populations (i.e., 15 ASVs), whether for cyanobacterial colony or FL fraction (Fig. 1C), revealing the prominent role of *Microcystis* as a contributor to the cyanopeptide pools. In comparison, *Dolichospermum*-affiliated ASVs were correlated with microginin FR6 and nodularin_R especially in the colony fraction (n = 2 links, *P* < 0.05), while *Cyanobium*-affiliated ASVs in cyanobacterial colony or PA fraction (n = 7 links, *P* < 0.05), reflecting differentiation in their respective interactions with cyanopeptides. Notably, all cyanopeptide components showed significant and positive inter-correlations between themselves (Pearson’s *r* range from 0.29 to 0.99, 0.68 on average, *P* < 0.05) (Fig. 1C, heatmap), revealing their co-occurrence during CyanoHABs in the lake Taihu.

### 3.2 Differentiation of non-cyanobacterial communities including microcystin degraders

Once cyanopeptides and their producers were identified, we further questioned on the corresponding dynamics of non-cyanobacterial communities and their ecological niches during CyanoHABs. Overall, the alpha diversity was greatly differed by size fraction with significantly lower diversity in the colony fractions, while the difference between sampling dates and lake regions are less prominent (Fig. S5). For beta-diversity, the non-cyanobacterial communities were also significantly and most differentiated by size fractions, followed by sampling dates (Fig. 3A & 3B). However, unlike cyanobacterial communities, their geographical heterogeneities (lake regions) were not statitically significant in explaining compositional variations (Table 1).

**Figure 3.**
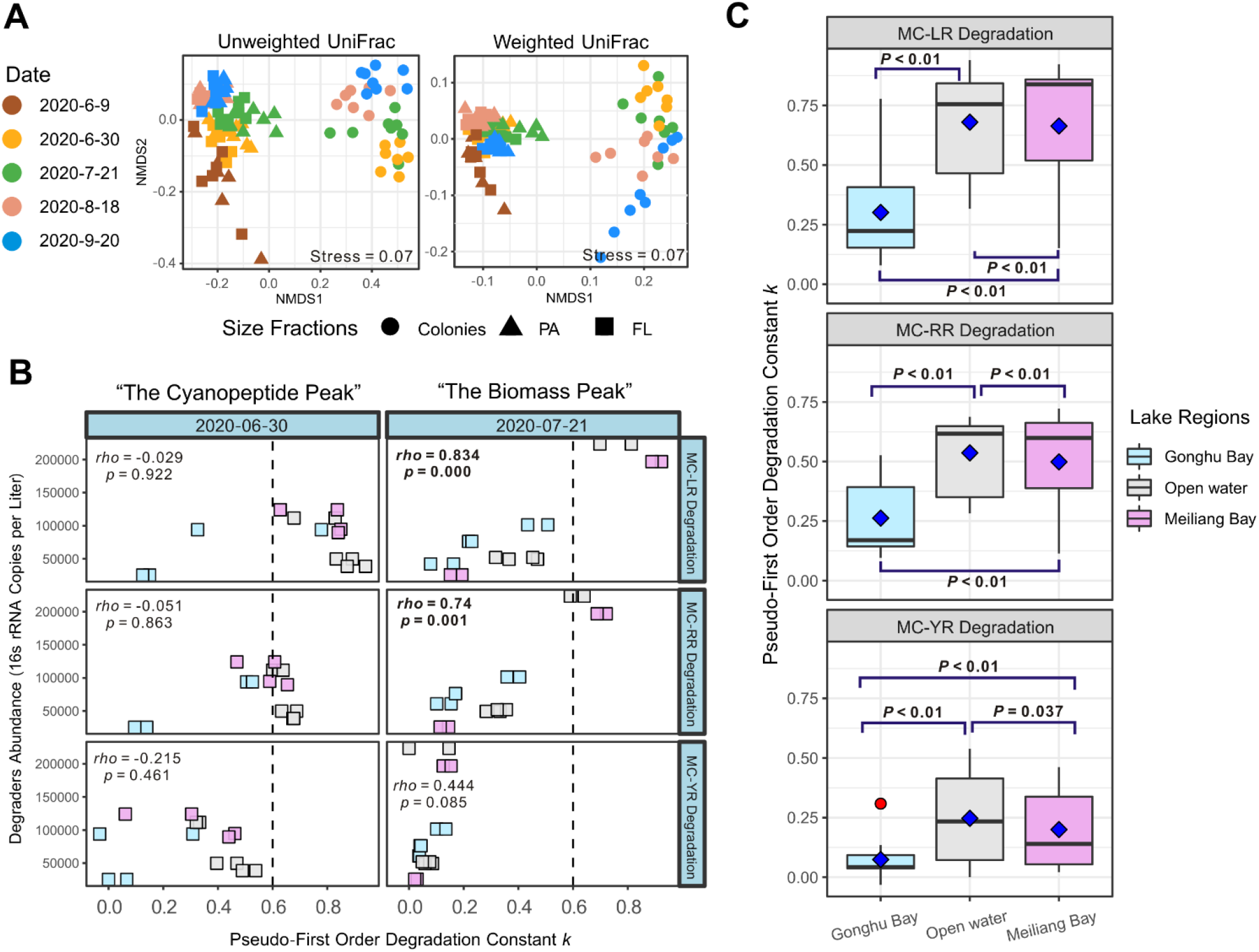
Spatiotemporal dynamics of non-cyanobacterial communities and cyanopeptides degraders in the Taihu Lake. (A) Non-multidimensional scaling (NMDS) analysis based on unweighted and weighted unifrac distances showing the community dissimilarity in terms of phylogenetic membership and abundance profiles, respectively. (B) Spearman’s rank correlations coefficients between the 16S-based absolute abundance of potential cyanopeptide degraders (copies L^-1^) and pseudo-first order degradation constant (*k*) values were calculated. (C) Boxplot showing degradation constant (*k*) from samples at separated sites as well as lake regions. Results revealed that the community composition was significantly differentiated by sampling date and size fraction. Abundance of potential degraders was positively correlated with degradation constant at the biomass peak (July 21) rather than cyanopeptide peak (June 30). Water samples from the center lake region (open water, Site 3, 4 and 8) exhibited significantly higher (T-test, *P* < 0.05) degradation capacity compared to the two bay areas.

Non-cyanobacterial communities are also known to harbor heterotrophs including cyanopeptides degraders. *Sphingopyxis* sp. was identified as the most prominent degrader in our lake water samples (Fig. S6, A, B). Using microcystins degradation kinetics (Fig. S7 and Fig. 3B), our microcosm experiments showed significant and strong positive correlations (*rho* > 0.74, *P* < 0.01) between degradation constant (*k*) and degraders abundance at the Biomass Peak on July 21, but contrastingly not at the Cyanopeptide Peak on June 30. Geographically, the results demonstrated that microbiota in the center lake region (open water, Site 3, 4 and 8) exhibited significantly higher (t-test, *P* < 0.05) degradation capacity compared to the two bay areas (Fig. 3C). Consistently, absolute abundance of *Sphingopyxis* sp. across the three lake regions decreased in the following order: Open water > Meiliang Bay > Gonghu Bay (Fig. S6C, inner bars), making the faster microcystin degradation in the open water than Meiliang and Gonghu Bay plausible.

### 3.3 Turnover of bloom-associated microbial assemblages ecologically determined by cyanopeptides

As cyanopeptides dynamics is shown to linked with the bloom-associated microbiota including their producers and degraders, we next question to which extent such deterministic selective forces contribute to the community assembly compared with stochastic factors (e.g., wind-driven mixing) and processes (i.e., dispersal and drift). Ecological stochasticity of microbial communities benchmarked by normalized stochasticity ratio (NST) showed that deterministic processes generally prevailed over stochastic processes (i.e., NST < 50%) in the colony fraction, particularly for cyanobacterial communities (NST = 22% to 40%, Fig. 4A), compared with non-cyanobacterial communities (NST = 46% to 55%, Fig. 4A). Intriguingly, cyanobacterial communities (Fig. 4A, top panel) and non-cyanobacterial communities (Fig. 4A, bottom panel) within PA and FL fractions displayed a distinct temporal dynamics of assembly patterns. For example, stochasticity of cyanobacterial community in PA and FL fraction was decoupled (Fig. 4A, top panel), reaching a peak on July 21 (Biomass Peak, NST = 98%) and Aug. 18 (NST = 99%), respectively. While the assembly of non-cyanobacterial communities was generally parallel, ascending until the Cyanopeptide Peak on June 30 (NST = 87% for PA and 86% for FL) and then declined (Fig. 4A, bottom panel).

**Figure 4.**
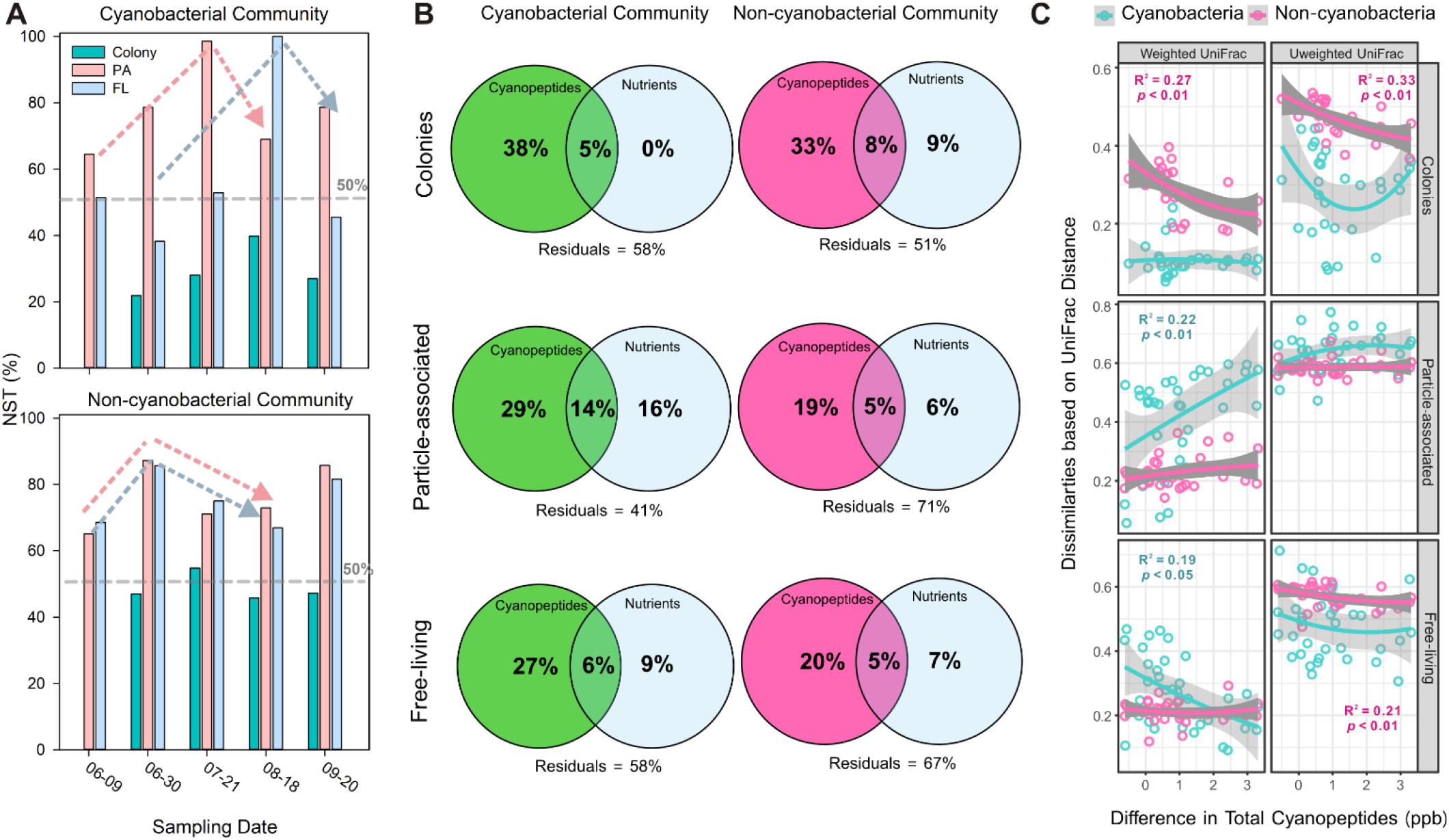
Cyanopeptides were identified as an important deterministic driver in turnover of both cyanobacterial and non-cyanobacterial communities. Comparison of NST (%) index of cyanobacterial (A, upper panel) and non-cyanobacterial communities (A, lower panel) in the three size-fractions. Colony samples on July 9 were not collected and therefore not shown. (B) Variation partitioning analysis (VPA) results show the relative contribution of cyanopeptides or nutrients to the size-fractioned cyanobacterial and non-cyanobacterial communities. (C) UniFrac distances of cyanobacterial/non-cyanobacterial communities, plotting against differences in total cyanopeptides concentrations. The lines were second-order polynomial regression curves fitted points of two communities denoted by colors. Only regression coefficients with *P* < 0.05 were annotated with a regression coefficient. Each point stands for a sample pair. The results generally showed deterministic effect of cyanopeptides on both cyanobacterial and non-cyanobacterial communities. And cyanopeptide was a factor stronger than nutrient in explaining turnover of bloom-associated microbiota. The community-level dissimilarities declined as a function of greater differences in cyanopeptide concentrations, with variances between size-fraction niches and communities.

To further discern the variation of the bloom-associated microbiota explained by deterministic factors, variation partitioning analysis (VPA) was employed (Fig. 4B). The results revealed that the nine cyanopeptides were much better in explaining the variances of cyanobacterial and non-cyanobacterial communities in all three size fractions, compared with the eight nutrient parameters (Fig. S2).

To quantitatively resolve the extents to which cyanopeptides dynamics in determining the assembly of bloom-associated microbiota, we sought for significant correlations between Chao1 richness and the nine cyanopeptide components (Fig. S8) and further fitted the dissimilarities (UniFrac distance) between bloom-associated communities against the concentration differences in total cyanopeptides (Fig. 4C). As expected, significant negative Spearman’s rank correlation coefficients (*rho*) were observed between Chao1 richness of non-cyanobacterial communities and cyanopeptides components (Fig. S8). The trend within cyanobacterial communities was similar, with negative *rho* values ranging from -0.244 to -0.445 for all nine cyanopeptide components (Fig. S8). On community-level, the influence of total cyanopeptide differences was the most evident in the colony fraction (Fig. 4C, top panels). Particularly, within the colony fraction, differences in cyanopeptide contents were negatively related with both the phylogenetic dissimilarity (R^2^ = 0.33, *P* < 0.01) and abundance-weighted dissimilarity (R^2^ = 0.27, *P* < 0.01) of those non-cyanobacterial assemblages (Fig. 4C). For communities in the PA and FL fractions, weighted UniFrac rather than unweighted UniFrac dissimilarities between cyanobacterial assemblages were significantly (*P* < 0.05) related to cyanopeptide variations but on opposite directions (Fig. 4C), revealing that cyanopeptide dynamics created distinct niches for cyanobacterial populations in the two size fractions.

## 4. Discussion

CyanoHABs across the globe have raised wide and ever-growing concerns on the eco-environmental and human health impacts of bioactive cyanopeptides (Janssen 2019). Here, for the first time, we expand current research scope on cyanopeptides beyond microcystins in Taihu and linked them to the size-fractionated lake water microbiota (i.e., cyanobacterial colony, particle-associated and free-living) over the whole course of a summer cyanobacterial bloom. High-resolution mass spectrometry and high-throughput DNA sequencing approaches provide new and complementary chemical and biological insights into those previously unconsidered but arguably harmful non-microcystin cyanobacterial metabolites, enabling quantitative elucidation of their interplay with microbiota and the key role of cyanopeptides in the microbiota assembly and ecosystem function underlying CyanoHABs.

### 4.1 Non-microcystin cyanopeptides of emerging ecotoxicological concerns

Limited knowledge on the occurrence and eco-impacts of cyanopeptides beyond microcystins hindered understanding on their toxicology and health risks (Janssen 2019, Natumi and Janssen 2020). In this study, we reported a comprehensive profile of cyanopeptides in lake Taihu including six major non-microcystin structures from four classes that were highly prevalent during a cyanobacterial bloom. The presence of non-microcystin cyanopeptides is not surprising, since their biosynthetic gene clusters in the dominant taxa have long been recognized (Calteau et al. 2014, Harke et al. 2016). What noteworthy is their high prevalence and compelling biological activities that make themselves emerging CyanoHABs-associated byproducts with deterministic but yet-under-appreciated eco-impacts (Swain et al. 2017).

Cyanostatin B, known as a potent inhibitor of leucine aminopeptidases (LAPs, EC3.4.11.1), was the most abundant cyanopeptide in our investigation (up to 1323.81 μg L^-1^). The concentration is 100 times higher than the reported IC_50_ (12 μg L^-1^) of LAP – M for microsomal from porcine kidney (Sano et al. 2005), while inhibition on microbial community are less reported. Considering the critical role of LAPs in microbial proteolysis and recycling of amino acids from peptides (Matsui et al. 2006), we speculated that the dominance of cyanostatin B as LAPs inhibitors during the lake bloom environment could hinder the biodegradation and further remineralization of cyanopeptides by heterotrophs. Consistent with our new hypothesis, potential inhibition of the cyanostatin B on the microbial richness were identified, as supported by the negative correlations with Chao1 richness index (Fig. S8). Likewise, nodularin_R is another putative potent inhibitor of microbial richness, given their negative correlations with both non-cyanobacterial and cyanobacterial communities (Spearman’s rank *rho* = -0.200 and -0.328, respectively). This cyanotoxin shares similar structure and inhibition potency as microcystins (e.g. IC_50_ of ∼2 nM or 1.65 μg L^-1^ for PP1), therefore may pose a synergistic toxic effect with microcystins (Honkanan et al. 1994, Huang and Zimba 2019). Moreover, although not been statistically detected as an inhibitor of microbial diversity, the concentration level of cyanopeptolin-1020 in the bloom waters (at a peak concentration of 7.1 μg L^-1^) is already high enough to exert toxicological effects on trypsin (IC_50_ = 670 pM or 0.68 μg L^-1^) of freshwater crustacean (Gademann et al. 2010) and zebra fish (Faltermann et al. 2014). Given more than 10 times higher concentrations of total non-microcystin cyanopeptides than the well-described microcystin detected in the lake Taihu, our results suggest that cyanopeptides produced during CyanoHABs should have been exerting hidden eco-impacts (e.g., microbial inhibition as afore-discussed) stronger than previously acknowledged microcystins alone. Therefore, future screening and profiling of cyanotoxins in lake ecosystems need to fully consider non-microcystin cyanopeptides as emerging ecotoxicologically critical components in governing aquatic biodiversity and explore their specific roles in mediating biogeochemical cycling in freshwater ecosystems plagued by CyanoHABs.

### 4.2 Degradation co-affected by putative cyanopeptide degraders and local microbiological setups

In natural water, the elimination of cyanobacterial secondary metabolites is revealed to mostly attribute to bacterial biodegradation (Li et al. 2017), which in turns involves in turnover of associated bacterial communities (Lezcano et al. 2017). Previous studies have evaluated the microcystin-degrading capacities of *Sphingopyxis* isolates from Taihu (Yan et al. 2012, Yang et al. 2020). Consistently, our study reveals that *Sphingopyxis* spp. are the most predominant microcystin degraders in lake water (Fig. S6). Moreover, we have experimentally verified the dependence of biodegradation capacity on the absolute abundance of the degraders at the Biomass Peak and also across three sampling regions. Interestingly, the positive ‘biomass-degradation’ relationship cannot be inspected at the Cyanopeptide Peak (on June 30) which came earlier than the Biomass Peak. This intriguing temporal disparity is likely due to cyanopeptides restriction on microbial diversity and degradation capacity, implying a dynamic interplay more complex than solely linear correlations between removal of cyanopeptides and biomass of degraders.

Field research has reported an antagonistic relationship between toxic cyanobacteria and microcystin degraders (Lezcano et al. 2018). Consider the inhibition effects of detected cyanopeptides as discussed in 4.1, we would presume that at least partial degrading microbes also be quantitively restricted to be decoupled from their respective cyanopeptide substrates, when cyanopeptides (thus their presumptive toxicities) peaked. Moreover, we can see that the non-linear relationship at Cyanopeptide Peak is featured by open water samples displaying high *k* values but relatively low degrader biomass (grey scatters, left panel), compared with the Biomass Peak (right panel, Fig. 3B). This result suggests that efficient degraders beyond we listed (Table S2) might be active when cyanopeptide concentration peaked. The degradation of cyanopeptides in natural water involves diverse microorganisms (Li et al. 2017, Zhang et al. 2020) and affected by multiple environmental variables (Maghsoudi et al. 2016, Moron-Lopez et al. 2017). While our results generally highlight the nutritional dependency of known degrading bacteria on cyanopeptides degradation, the role of other unknown degrading processes and cross-feeding of degrading products (e.g., phenylacetate) cannot be ruled out. For instance, the peaked cyanopeptides can create a hotspot of organic carbon (Su et al. 2018, Zhang et al. 2020) to stimulate microbial degradation via pathways alternative to those in *Sphingopyxis* cells (Lezcano et al. 2017, Moron-Lopez et al. 2017), resulting in the high degradation rates at low biomass of reported degraders as detected in the open water (grey scatters, left panel, Fig. 3B). In addition, degradation of cyanopeptides would be favored in indigenous biological setups where pre-adaption (Christoffersen et al. 2002, Cousins et al. 1996, Moron-Lopez et al. 2017) and quorum sensing system (Zeng et al. 2020) can both extensively regulate and enhance microbial degradation capacity. Therefore, the lower *k* values observed for sites within Meiliang bay and Gonghu Bay (Fig. 3C) could result from the premature microbiological setups for cyanopeptide degradation, as the two bay area are more likely be subject to influence of water diversion and inflow than the open water regions (Hu et al. 2021, Zhu et al. 2021b).

Collectively, our results indicate degradation of cyanopeptides in the lake Taihu is co-determined by abundance of putative degraders and local microbiological setups. Geographically, the increased water inflow (by 38.5%, i.e. around 3 billion m^3^ per year) and accelerated water exchange cycle (from 210 days to 184 days) since 2007 (Zhu et al. 2021b) could also interfere the remediation and restoration (in this case, biodegradation of the cyanopeptide during bloom event) by indigenous populations in Taihu, implying a trade-off between geo-engineering and ecosystem resilience. It should be noted that the above speculation was made by extrapolating degradation kinetics of microcystins to all cyanopeptides, while for a group of cyclic or linear peptides with conserved substructures in each class (Janssen 2019), variances in degradation efficiency are expected and deserve further investigation.

### 4.3 Cyanopeptides as deterministic factors of bloom-associated microbiota assemblage

Recent studies have suggested that the turnover of microbial communities is tightly associated with CyanoHABs due to prevalent nutrient cycling and signal transduction between bloom-associated microbiota (Wang et al. 2020, Zhu et al. 2021a, Zhu et al. 2019), as well as dynamic of microcystins (Lezcano et al. 2017). Here, we extended the scope to cyanopeptide components beyond microcystin and showed that assembly of bloom-associated microbiota is strongly determined by cyanopeptides, with variances between size-fraction niches and communities.

Cyanobacterial colony represents a unique niche for bloom microbiota where cyanopeptides regulate both cyanobacterial and non-cyanobacterial populations. Sizes of colonies are positively correlated with microcystin concentrations (Gan et al. 2012, Wang et al. 2013), implying an intrinsic connection between cyanopeptides and cyanobacterial community. This is consistent with our observation that deterministic processes mostly contributed to the assembly of cyanobacterial communities within colonies (Fig. 4A). The intriguing co-existence of multiple closely related populations (as proxied by ASVs) and their changing abundances were observed for cyanobacterial genera in the colony (Fig. 2C), reflecting a combined role of cyanopeptide dynamics and environmental filtering in selecting competing populations and driving community turnover. Meanwhile, assembly of non-cyanobacterial communities within the colony was basically controlled by deterministic processes except for that during the Biomass Peak (Fig. 4A, bottom panel), and the community was also more responsive to cyanopeptides dynamics than nutrients (Fig. 4B). Moreover, their community-level dissimilarities within colonies declined as a function of larger differences in cyanopeptide concentrations (Fig. 4C), indicating a much narrower range of ecological niches (thus lower observed alpha diversity, Fig. S4 & S5) for the colonized microbiota. Similarly, the fit of bacterioplankton communities to neutral model was weaken due to biotic pressure from accompanied cyanobacteria (Wang et al. 2020). Here, our results further indicate that cyanopeptides including those bioactive non-microcystin ones (as afore-discussed in 4.1), could be chemical intermediates of the biotic pressure. Colonies also nurture the most abundant *Bacteroidetes* at the Biomass Peak (Fig. S3A). This phylum members can readily adapt to complex substrates rather than monomeric compounds and therefore play a fundamental role in conversion of high molecular weight planktonic compounds during phytoplankton blooms (Buchan et al. 2014). In this scenario, the increased stochasticity at the Biomass Peak (July 21) could be plausibly co-explained by the rapid degradation of cyanopeptides (from ca. 597 to 300 ppb on average) into labile products (e.g. amino acid) to nourish heterotrophic communities (as discussed in 4.2), and also the relief from the restrictive pressure, as noticed elsewhere (Zhou et al. 2014).

For PA and FL microbiota, the distinct dynamic of assembly patterns for cyanobacterial and non-cyanobacterial communities, varied contributions of cyanopeptides or nutrients in explaining community turnover and the contrasting community-level responses to cyanopeptides (Fig. 4) together suggest that assembly of the two communities was dominated by separated processes throughout the bloom event. For example, the increased in stochasticity for cyanobacteria in PA fraction from June 09 to July 21 (Fig. 4A, top panel) could originate from the expansion of *Microcystis* spp. in the PA metacommunity (Fig. 2B, PA fraction), which is more likely to be taxonomically closer to *Microcystis* populations in the colony (Zhu et al. 2018), leading to a random distribution pattern and in turn, high stochasticity (Ning et al 2019). Cyanopeptides provide their producers competitive edge over associated populations (Holland and Kinnear 2013). As the free-living cyanobacteria are phylogenetically more diverse than communities in the colony and PA fraction, high cyanopeptide levels may have mediated their stochasticity until the producers were less prominent in biomass (i.e. on Aug. 18). This may have also explained the opposite trend of cyanobacterial dissimilarities in PA and FL fraction (Fig. 4C), as competition exclusion induced by cyanopeptides drived the FL communities more phylogenetically similar (Chen et al. 2020, Mayfield and Levine 2010). Collectively, our results highlight that cyanopeptides, through their broad microbial restriction effects as well as degradation, are the key biotic forces in mediating microbiota assembly during CyanoHABs.

## 5. Conclusion

Temporal and spatial dynamics of cyanopeptides in the lake Taihu during summer CyanoHABs were closely followed and linked to the microbiota to elucidate its assembly mechanisms. Cyanopeptides represent a variety of structural classes and can exert significant ecotoxicological impacts on bloom-associated microbiota. Biomass of potential degrading microbes could quantitatively indicate the degradation capacity of lake water, while the spatiotemporal disparity suggested a complex interplay between removal of cyanopeptides and local microbiota setup which is subject to influences of water division and inflow. Furthermore, cyanopeptide was identified as a selective force (stronger than nutrients) in mediating the assembly and turnover of bloom-associated microbiota, with variances between size-fraction niches and communities. Overall, we propose that the prevalence, dynamics and eco-impacts of cyanopeptides should be considered as a crucial consequence of intractable CyanoHABs and deserve oriented surveillance and investigation in future to address their ecological roles in ecosystems like Taihu.

## Acknowledgment

This work was supported by the Young Scientists Fund of the National Natural Science Foundation of China (Grant No. 51908467, F.J.) and Institutional Funds from Westlake University and Westlake Institute for Advanced Study (F.J.). We thank Prof. Wei Zhu and his group members at Hohai University for their great support in sampling and nutrients analysis. We thank Mr. Jinheng Pan and Ms. Xiuxiu Zhao from Mass and Metabolism Facility at Westlake University for their assistance with mass spectrometry.

